# Elucidation of dose-dependent transcriptional events immediately following ionizing radiation exposure

**DOI:** 10.1101/207951

**Authors:** Eric C. Rouchka, Robert M. Flight, Brigitte H. Fasciotto, Rosendo Estrada, John W. Eaton, Phani K. Patibandla, Sabine J. Waigel, Dazhuo Li, John K. Kirtley, Palaniappan Sethu, Robert S. Keynton

## Abstract

Long duration space missions expose astronauts to ionizing radiation events associated with highly energetic and charged heavy particles. Such exposure can result in chromosomal aberrations increasing the likelihood of the development of cancer. Early detection and mitigation of these events is critical in providing positive outcomes. In order to aid in the development of portable devices used to measure radiation exposure, we constructed a genome-wide screen to detect transcriptional changes in peripheral blood lymphocytes shortly after (approximately 1 hour) radiation exposure at low (0.3 Gy), medium (1.5 Gy) and high (3.0 Gy) doses compared to control (0.0 Gy) using Affymetrix^®^ Human Gene 1.0 ST v1 microarrays. Our results indicate a number of sensitive and specific transcriptional profiles induced by radiation exposure that can potentially be implemented as biomarkers for radiation exposure as well as dose effect. For overall immediate radiation exposure, *KDELC1*, *MRPS30*, *RARS*, and *HEXIM1* were determined to be effective biomarkers while *PRDM9*, *CHST4*, and *SLC26A10* were determined to be biomarkers specific to 0.3 Gy exposure; *RPH*, *CCDC96*, *WDYHV1*, and *IFNA16* were identified for 1.5 Gy exposure; and *CWC15*, *CHCHD7*, and *DNAAF2* were determined to be sensitive and specific to 3.0 Gy exposure. The resulting raw and analyzed data are publicly available through NCBI's Gene Expression Ominibus via accession GSE64375.

## Introduction

The National Aeronautics and Space Administration Authorization Act of 2010 (1) and the National Space Policy of the United States of America (2010) (2) set in motion the goals for cislunar and deep space exploration. Among these goals are manned missions for the Asteroid Redirect Mission (3) and Journey to Mars (4). The long duration of these missions (up to 1,100 days), necessitates the development of lightweight and portable devices for monitoring health. The top health risk for astronauts on such missions beyond low earth orbit (LEO) is exposure to ionizing radiation associated with highly energetic and charged heavy particles originating from constant galactic cosmic rays and sporadic solar particle events (5–7). This exposure can lead to increased risk of cancer (8–14); deficits in the central nervous system (15–23); degenerative tissue effects (24, 25) including changes attributed to the increase in oxidative stress that are normally associated with aging, such as cataract formation (26–28) and vascular degeneration (29, 30); and acute radiation syndrome (31–34) marked by decreased circulating blood cells (35, 36), lung damage (37), decreased cardiac function (38, 39), and immune system suppression (5).

Current methodologies for measuring radiation exposure still have a high degree of uncertainty when it comes to determining the level of radiation to which an astronaut crew member has been exposed (5, 40–42). Most of the methods to date have focused on detecting chromosomal aberrations, including single-(SSB) and double-stranded (DSB) breaks, translocations, and exchanges. DSBs have been used extensively, due to their direct relation with radio-induced biological effects, including unequal crossover, chromosomal rings (43), inversions, and dicentric chromosomes (44–46). Detection of these aberrations using either Giemsa staining techniques or fluorescent in-situ hybridization (FISH) (7, 47) have become the *de facto* gold standard for biological dosimetry, showing a dosage exposure accuracy rate of ± 10% (48). While these are currently the most reliable methods in detecting chromosomal aberrations, it is likely the development of high-throughput genome-wide screens of radiation exposure will allow for the detection of sensitive and specific transcriptional biomarkers which can be used in biodosimeters for detecting changes in response to radiation (47).

On an individual gene level, a number of genes have been used as biomarkers for radiation exposure at either the transcript, protein, or modified protein level, including phosphorylated H2A Histone Family, Member X (γ*H2AFX*), Tumor Protein 53 (*TP53*), and Cyclin-Dependent Kinase Inhibitor 1A (*CDKN1A*). Detection of the phosphorylation of *H2AFX* has been used in assays to determine radiation exposure due to its role in DNA double-stranded break repair (49–53) and is perhaps the best example of a single protein modification used for radiation detection. *TP53* is known to function as a transcription factor, which is radiation-modulated (54–59), and *CDKN1A* is a downstream target of *TP53*, which regulates progression through the cell cycle (60–62).

While each of these biomarkers have been shown to be sensitive to radiation exposure, they fall short in their specificity at a transcriptional level. Therefore, the purpose of this work is to identify additional transcriptional biomarkers which are both sensitive and specific to low, medium, and high levels of radiation at 1 hr post-exposure. The focus of our study is specifically on ionizing radiation from γ-rays and does not consider the specific effects of other sources of ionizing radiation (such as α-particles, β-particles, and positrons) which may also have an effect on transcriptional activity.

## Materials and methods

### Experimental design

Details of the study design have been previous described in Rouchka et al. (63). In summary, the design was constructed to determine transcriptional changes within white blood cells (WBC) at an early time point (1 hr) following ionizing radiation exposure. Whole blood was drawn from four volunteers prior to irradiation. Samples were maintained at room temperature throughout the radiation and WBC isolation process. The blood samples were then irradiated with γ-radiation using a Gammacell 1000 Elite (Noridon) for 0s (0.0 Gy – control), 3s (0.3 Gy), 16s (1.5 Gy), or 32s (3.0 Gy) with four biological replicates at each level (n=4). After completion of the radiation cycle, WBC were separated according to the methods in (63). Total RNA was then extracted from the WBC for use in the microarray experiments. All procedures were done in accordance with published NASA and NIH Guidelines, the University of Louisville Institutional Review Board (IRB), and the University of Louisville Institutional Biosafety Committee (IBC).

### Microarray preparation

Microarrays were prepared using Affymetrix^®^ Human Gene 1.0 ST v1 microarrays (Gene Expression Omnibus (GEO) (64) platform GPL6244 transcript version; GPL10739 probeset version). After the samples were prepared and the biotinylated cDNA was hybridized to the arrays, microarrays were scanned using an Affymetrix^®^ GeneChip^®^ Scanner 3000 and the GeneChip^®^ Command Console^®^ software version 3.1 (Affymetrix^®^) resulting in 16 raw CEL files (Table 1).

**Table 1.**
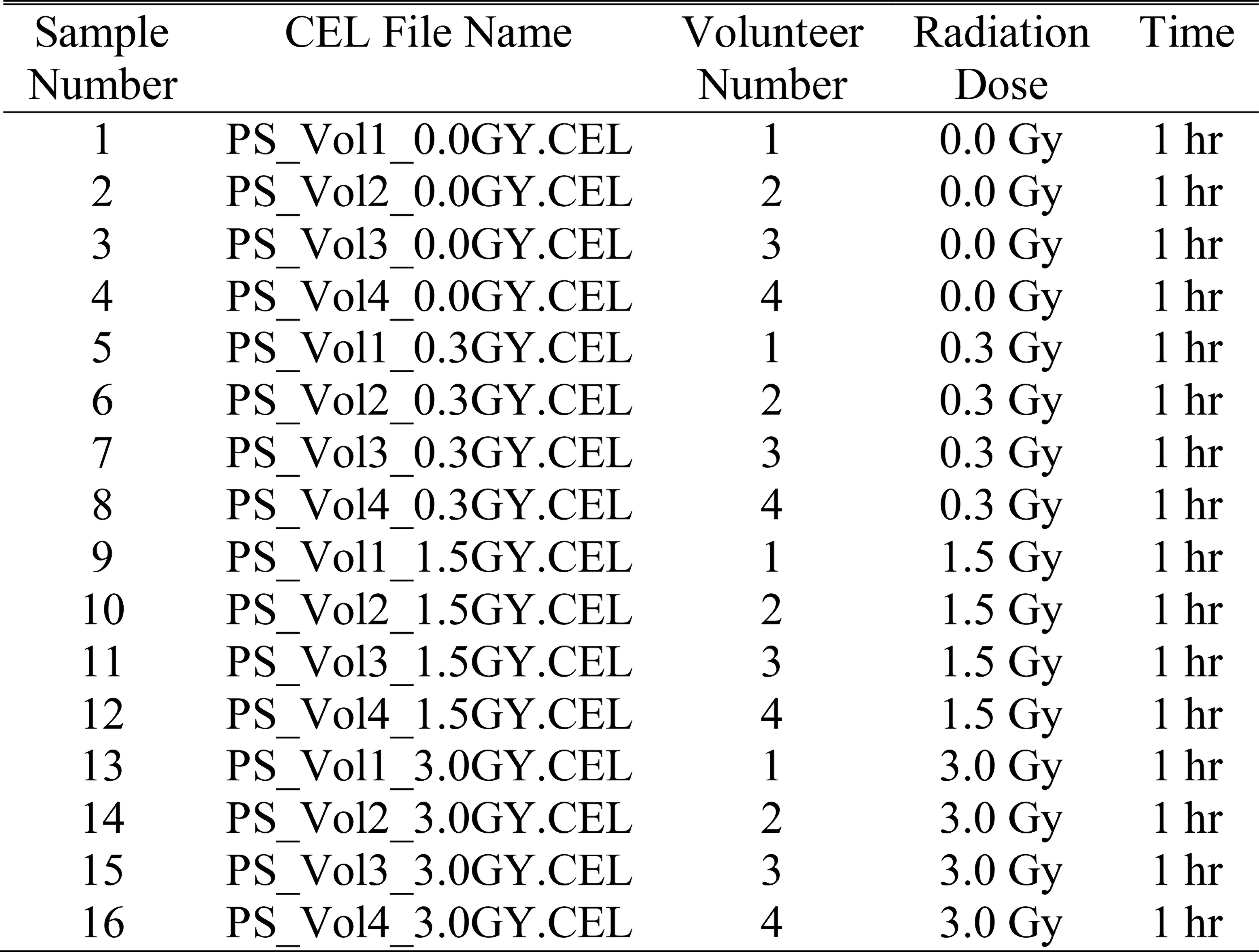
Microarray samples.

### Microarray analysis

Raw microarray files were processed in RStudio (65) (version 0.98.501) using R (66) (version 3.0.1 2013-05-16 “Good Sport”) and packages obtained from CRAN (67) and Bioconductor (68). CEL files were preprocessed and normalized using the oligo package (69) and robust multichip averaging (RMA) (70) based on the default Affymetrix^®^ transcript chip description format (CDF) (GEO platform GPL6244) which organizes the probes into 33,297 transcripts. CEL files were organized into four phenotypic categories according to the descriptions in Table 1, including four biological replicates at each of the four radiation doses of 0.0 Gy (control), 0.3 Gy, 1.5 Gy, and 3.0 Gy. A single contrast matrix was constructed in order to complete three dose-dependent comparisons: 0.3 Gy (n=4) vs. control (n=4); 1.5 Gy (n=4) vs. control; and 3.0 Gy (n=4) vs. control. Note that in each comparison, the control remains the same untreated samples for the four volunteers (n=4). Differentially expressed genes (DEGs) (defined as Affymetrix^®^ transcript sets) were determined using the Limma (71) linear modeling Bioconductor package.

### Categorical enrichment

Entrez identifiers for differentially expressed probe sets were used as inputs into category Compare (72) for further analysis of enriched Gene Ontology Biological Processes (GO::BP)(73) and KEGG Pathways (74). For the GO::BP categories, a minimum gene count of 5 was used along with a p-value cutoff of 0.01, which was calculated using a hypergeometic test.

### Sensitive and Specific Transcriptional Biomarkers

In order to account for specificity, we analyzed publicly available GEO DataSets, which were highly curated and standardized experiments contained within GEO. As of 12/13/2015, a total of 3,848 GEO DataSets existed, out of a total of 63,462 GEO Series. Using the GEO DataSets as a standard, the corresponding gene symbols for the differentially expressed probe sets from our microarray study were searched against GEO DataSets to filter common genes that were not frequently up- or down-regulated. Filtering by commonly measured genes removed nearly all of the noncoding RNAs (snRNA, snoRNA, miRNA, lncRNA), since they were not commonly represented on all microarray platforms. For our purposes, a gene was determined to be common if it was found in at least 2,000 of the GEO DataSets, and infrequently differentially expressed if it was up- or down-regulated in fewer than 100 of the datasets. We further reduced the analysis to only up-regulated genes from our dataset, since the primary goal of our study was to identify biomarkers for developing technologies that would work best on determining detectable levels of gene products.

## Results

### Differentially expressed probesets

Using a p-value cutoff of 0.05, a total of 439 probesets were determined to be differentially expressed at 0.3 Gy vs. control; while 403 were found to be differentially expressed at 1.5 Gy and 494 at 3.0 Gy out of a total of 28,869 non-control probesets (Table 2). Among the top 10 transcripts differentially expressed at 0.3 Gy were *DPP10*, *KDELC1*, *MAGEA10*, *MS4A5*, *OR8I2*, *WNT5A* which were up-regulated and *CXCL11*, *OR1F2P*, and *STRA8* which were down-regulated. At 1.5 Gy, the top 10 differentially expressed transcripts included the up-regulated transcripts *KDELC1*, *METTL2A*, and *PCNA* and down-regulated transcripts *HIST2H2BE*, *HSPB8*, *OR4M2*, *PRR32*, *ROPN1*, and *STRA8*. The top transcripts differentially expressed at 3.0 Gy exposure included *HEXIM1*, *KDELC1*, *OR8I2*, *PCNA*, and *ZNF22* which were up-regulated, and *FAM47A*, *HIST2H2BE*, *HSD11B2*, *KLHDC9*, and *OR4M2* which were down-regulated. Expression patterns for the top 10 differentially expressed transcripts at each exposure level are given in Fig 1.

**Table 2.**
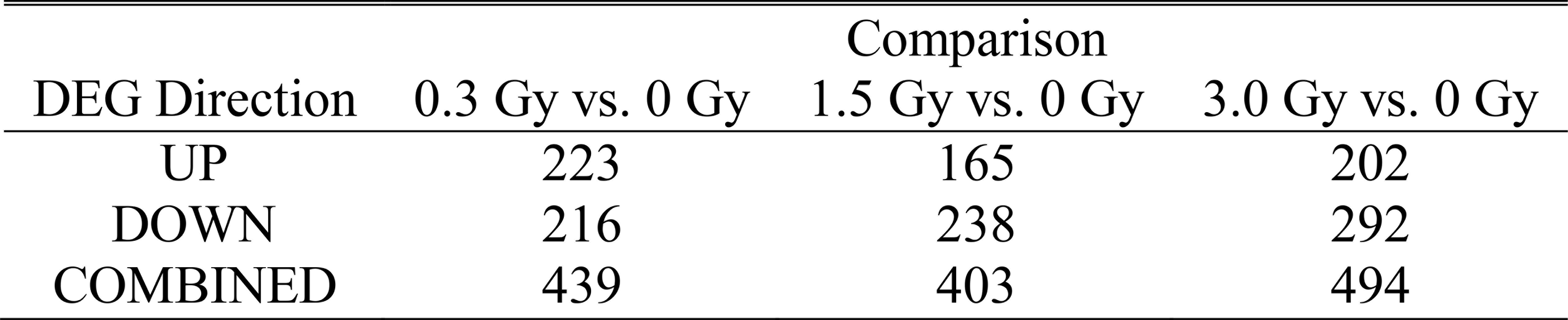
Number of differentially expressed transcript probesets for radiation dependent results (p-value ≤ 0.05; 1 hr post-exposure).

**Fig 1.**
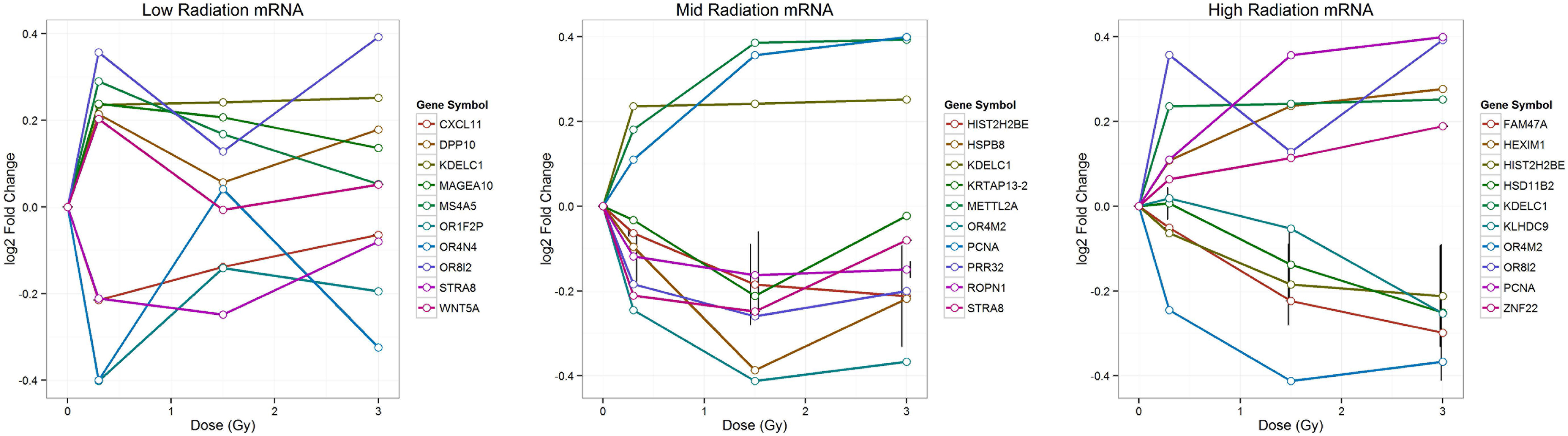
Gene expression patterns for top 10 differentially expressed transcripts at 0.3 Gy (left), 1.5 Gy (center) and 3.0 Gy (right).

Among the top differentially expressed transcripts that were up-regulated, *KDELC1* (KDEL motif-containing 1) was shared at all radiation levels. *KDELC1* is an endosplasmic reticulum protein. The function of the KDEL motif is to prevent all endoplasmic reticulum resident proteins from being secreted. *PCNA* (proliferating cell nuclear antigen) was among the top 10 differentially up-regulated genes at 1.5 and 3.0 Gy. Due to the higher level of radiation exposure, the up-regulation of *PCNA* was likely functioning in DNA repair through the *RAD6* pathway.

Shared differentially expressed down-regulated transcripts included *STRA8* (stimulated by retinoic acid) and *HIST2H2BE* (histone cluster 2, H2be). *STRA8* is thought to play a role in spermatogenesis. The down regulation of *STRA8* along with *SPATA2* and *ROPN1* could help to potentially reduce germ cell mutagenesis induced by radiation exposure (75). As a histone core protein, the down-regulation of *HIST2H2BE* may be related to the activity of one of its binding partners, *H2AFX* and likely plays a role in DNA accessibility and repair.

Two olfactory receptors, *OR8I2* and *OR4M2* were shared as differentially expressed transcripts. In the first case, *OR8I2* was up-regulated, while in the second case, *OR4M2* was down-regulated. The differential expression of olfactory receptors is not surprising, given the association of changes in sensory perception with radiation exposure (76–79).

Fig 2 shows a Venn diagram of shared gene symbols (based on Entrez (80) identifier) across the three radiation levels, with those shared between all three sets given in Table 3. As Table 3 illustrates, a number of the common transcript probes belong to small non-coding snRNA and snoRNA, which are involved in rRNA maturation. In order to illustrate that the direction of change was rather consistent no matter the radiation level, a heatmap was constructed to show the log_2_fold-change for all probes where the absolute value of the fold change was greater than or equal to 1.2 in at least one of the radiation exposure datasets (Fig 3).

**Fig 2.**
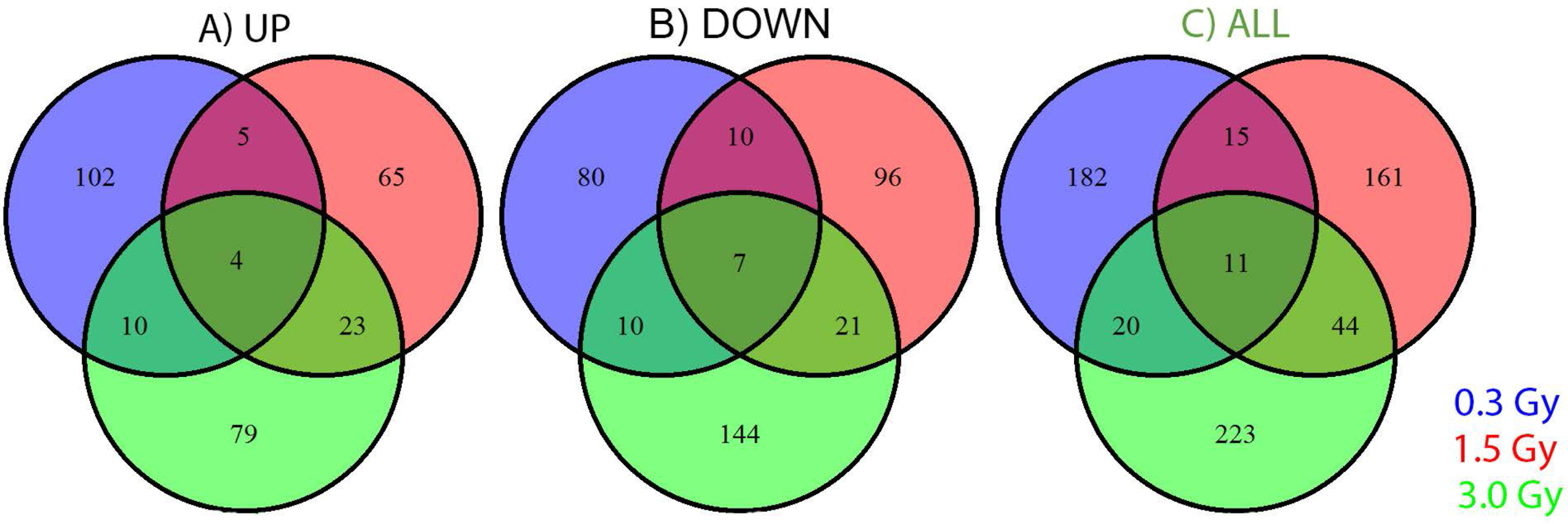
Venn diagram of shared gene symbols across radiation levels. Shown are the number of symbols that are A) up-regulated; B) down-regulated; C) up- and/or down-regulated. Additional probes without corresponding Entrez gene identifiers or gene symbols are not shown.

**Table 3.**
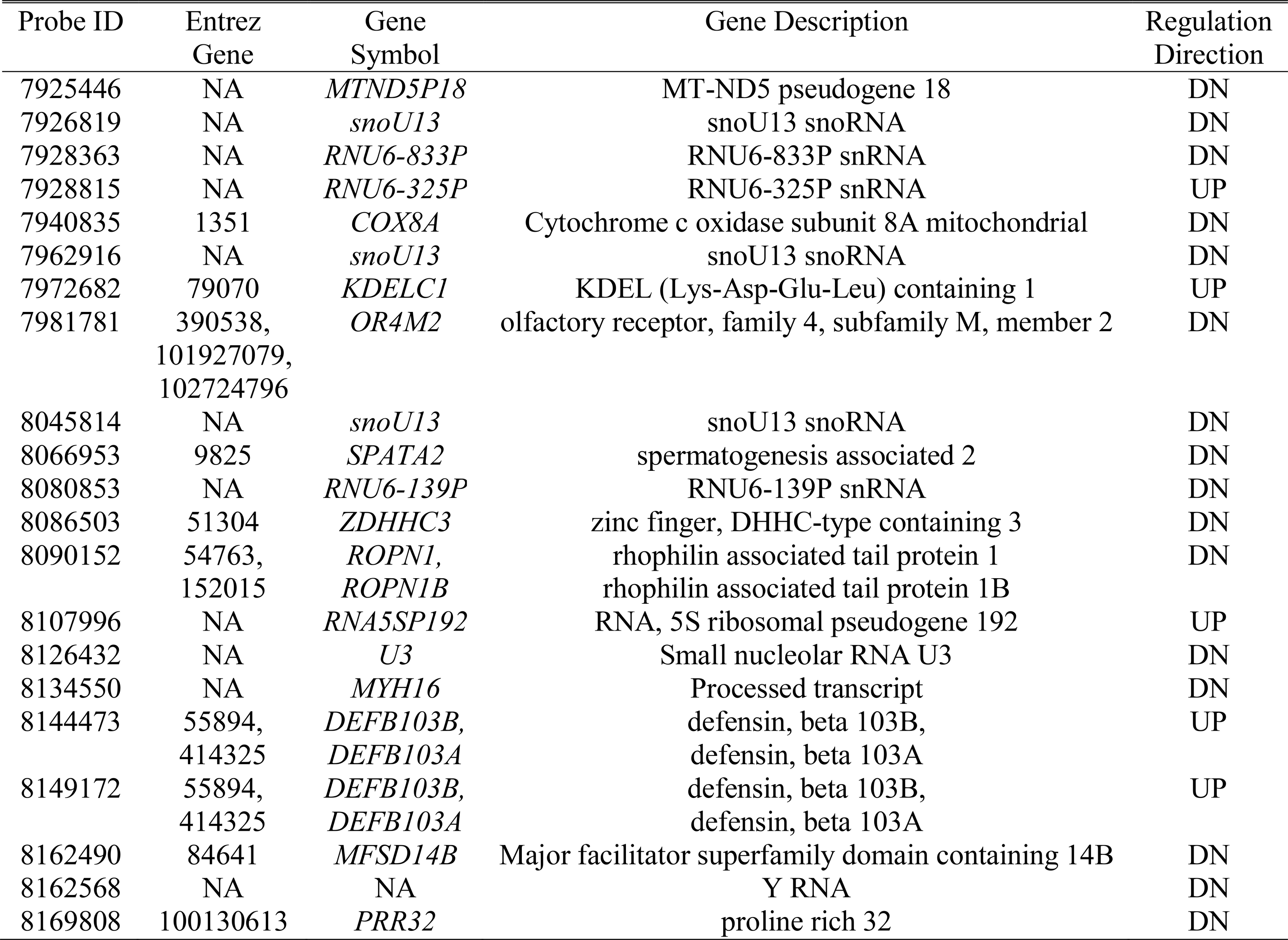
Common differentially expressed transcript probesets across all radiation levels.

**Fig 3.**
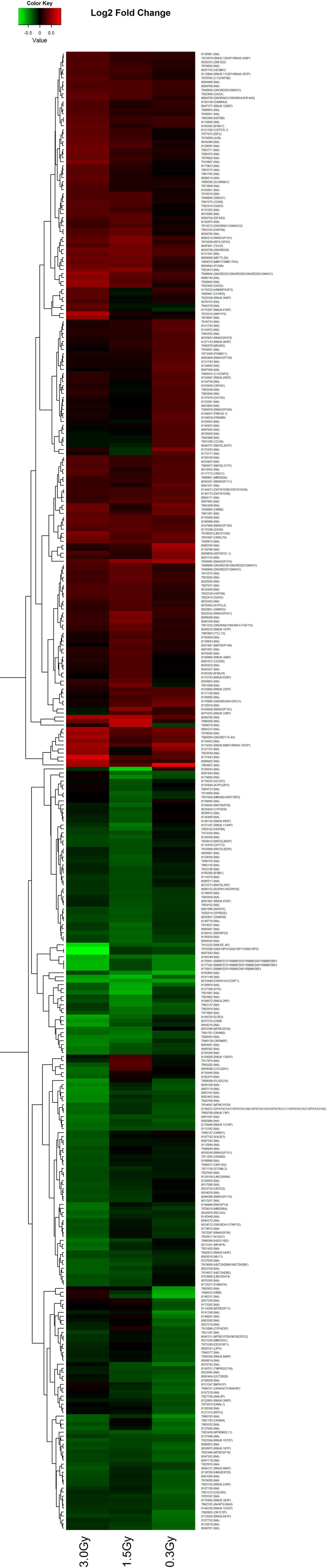
Heatmap of genes with at least one probe having FC of ±1.2 in at least one radiation level for the given condition in treated vs. control. Up-regulated genes are in red; down-regulated in green. Shown are radiation dependent results (three radiation levels vs. no radiation at time 1 hr).

### Differentially expressed microRNAs

In addition to protein coding genes, a small number of non-coding microRNAs (miRNAs) were found to be differentially expressed (Table 4; Fig 4). Among the differentially expressed miRNAs at 0.3 Gy radiation level, *MIR200A* has been shown to play a role in controlling epithelial-to-mesenchymal transition (EMT) (81, 82) as well as controlling oxidative stress response (83) and *MIN181B1*, which was down-regulated, modulates expression of *PTEN* (phosphatase and tension homolog) (84), a tumor suppressing gene mutated in a multitude of cancers. *MIR191* helps to trigger keratinocyte senescence by down-regulating the cell cycle regulator *CDK6* (cyclin-dependent kinase 6) (85). The miRNA *MIR520C* has been implicated in reduction of protein translation and induces cell senescence (86). Little is known about the function of the additional significantly differentially expressed miRNAs *MIR194-1*, *MIR323A*, and *MIR519A1*.

**Table 4.**
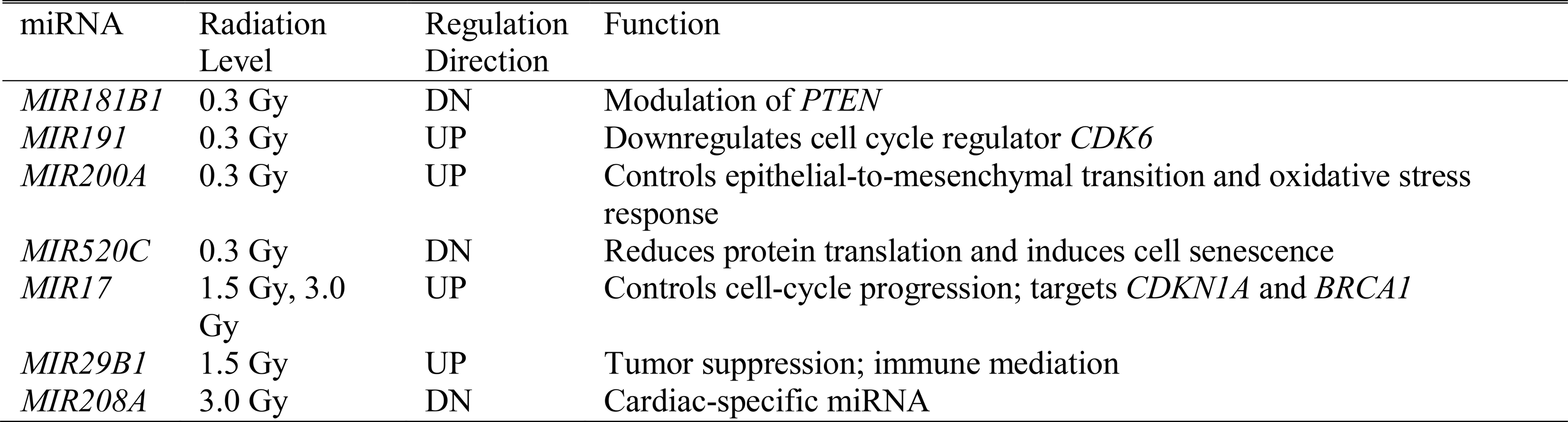
Differentially expressed miRNAs with known function.

**Fig 4.**
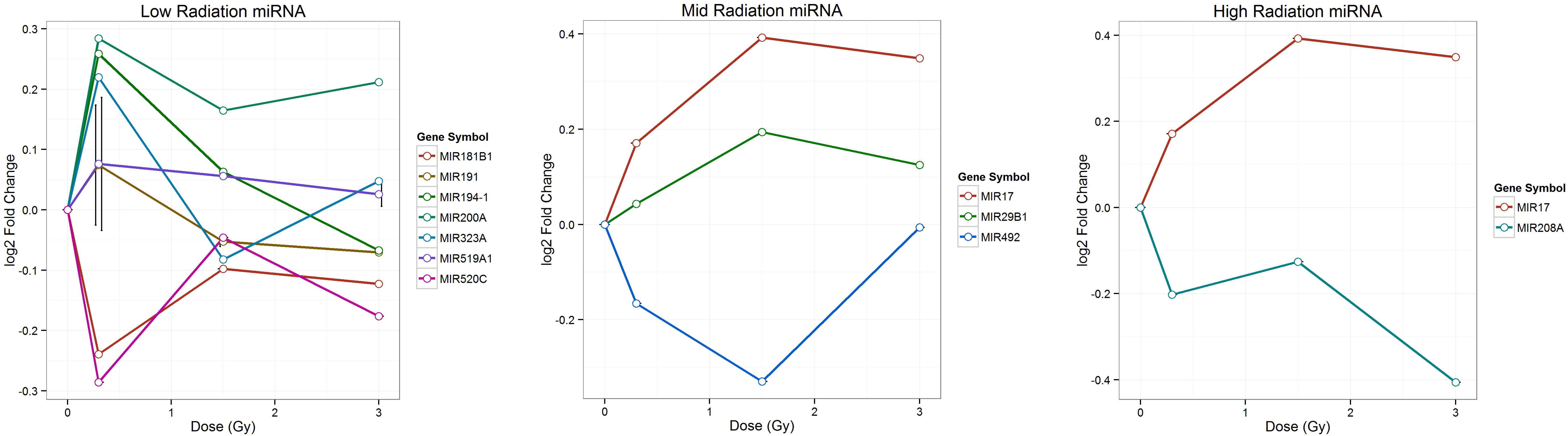
MicroRNA (miRNA) expression patterns for differentially expressed miRNA transcripts at 0.3 Gy (left), 1.5 Gy (center) and 3.0 Gy (right).

At 1.5 Gy, *MIR29B1* has been shown to have a tumor suppressor role through its control of methylation patterns by modulating the DNA methyltransferases *DNMT3A* and *DNMT3B* (87), its control of *CD276* (88), and regulation of a number of genes involved in acute myeloid leukemia (89). In addition, *MIR29B1* helped control the anti-apoptotic protein *MCL1* (myeloid cell leukemia 1) (90) and was down-regulated by the transcription factor *MYC* (91) as well as served a role in immune mediation (92). Little was known about the function of *MIR492*.

The miRNA *MIR17* was significantly up-regulated at both 1.5 Gy and 3.0 Gy. It has been shown to be activated by *MYC* and negatively regulated the *E2F1* transcription factor (93), thus providing a control mechanism for cell cycle progression. By controlling *E2F1*, *MIR17* helped to limit double-stranded breaks (94) and targets *CDKN1A* (cyclin-dependent kinase inhibitor 1A; P21) (95) and *BRCA1* (96) and, thus, was likely trying to recover from the effects of the damage induced by the radiation exposure. *MIR208A*, which was significantly down-regulated at 3.0 Gy, has been shown to serve as a cardiac-specific miRNA by controlling expression of the cardiac muscle myosin heavy chain 7b, *MYH7B* (97).

### Differentially expressed ncRNAs

In addition to miRNAs, a number of non-coding RNAs (ncRNAs) were determined to be significantly differentially expressed, particularly at higher radiation levels (Table 5; Fig 5). Among these were long, non-coding RNAs (lncRNAs), small nuclear RNAs (snRNAs), and small nucleolar RNAs (snoRNAs). The snRNAs and snoRNAs are largely functional in ribosomal RNA maturation.

**Table 5.**
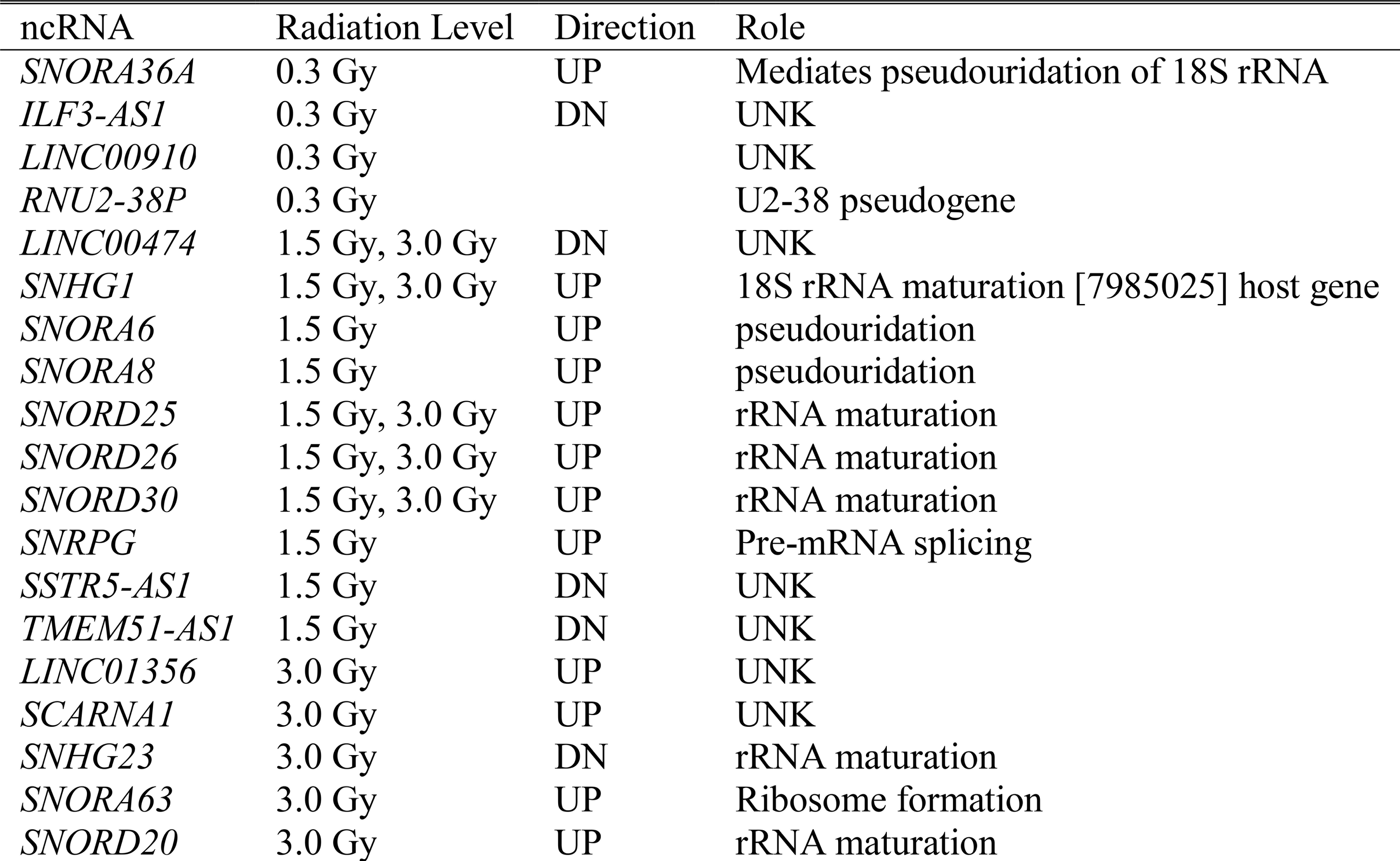
Differentially expressed ncRNAs and their associated function.

**Fig 5.**
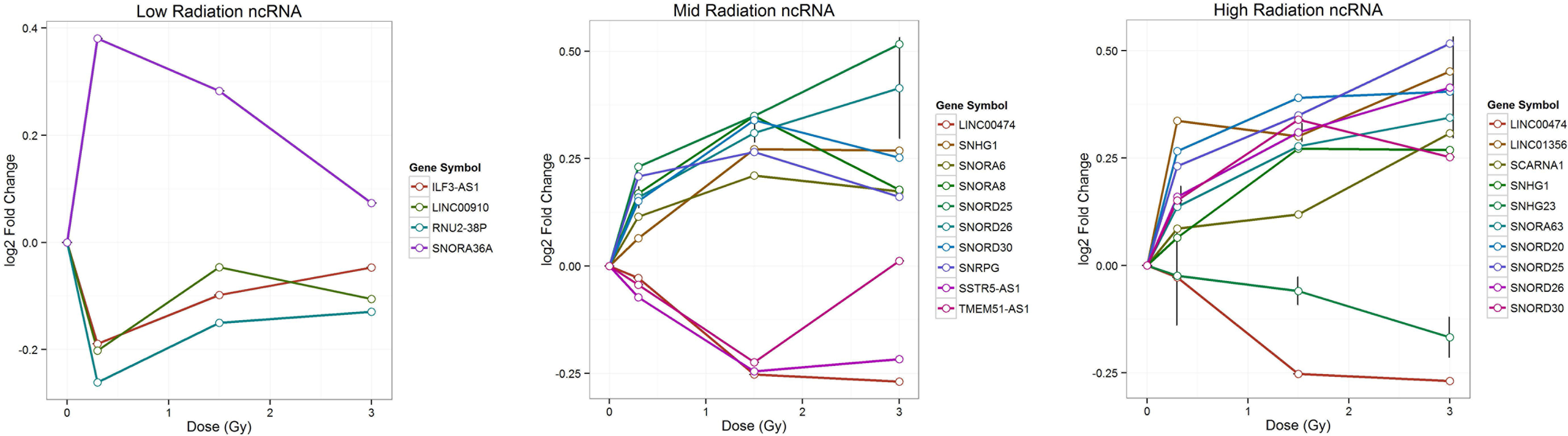
Non-coding RNA (ncRNA) expression patterns for differentially expressed ncRNA transcripts at 0.3 Gy (left), 1.5 Gy (center) and 3.0 Gy (right).

### Differentially expressed pseudogenes

Interestingly, a number of the significant differentially expressed probes belonged to pseudogenes for both protein coding and non-coding RNAs (Fig 6). Among the up-regulated pseudogenes were a number of ribosomal associated pseudogenes; two that were associated with olfaction (*VN2R17P*, *OR2U1P*); and one, *GCOM2* (*GRINC1B* complex locus 2, pseudogene) that was implicated in both myeloid leukemia (98) and telomeric repeat binding (99).

**Fig 6.**
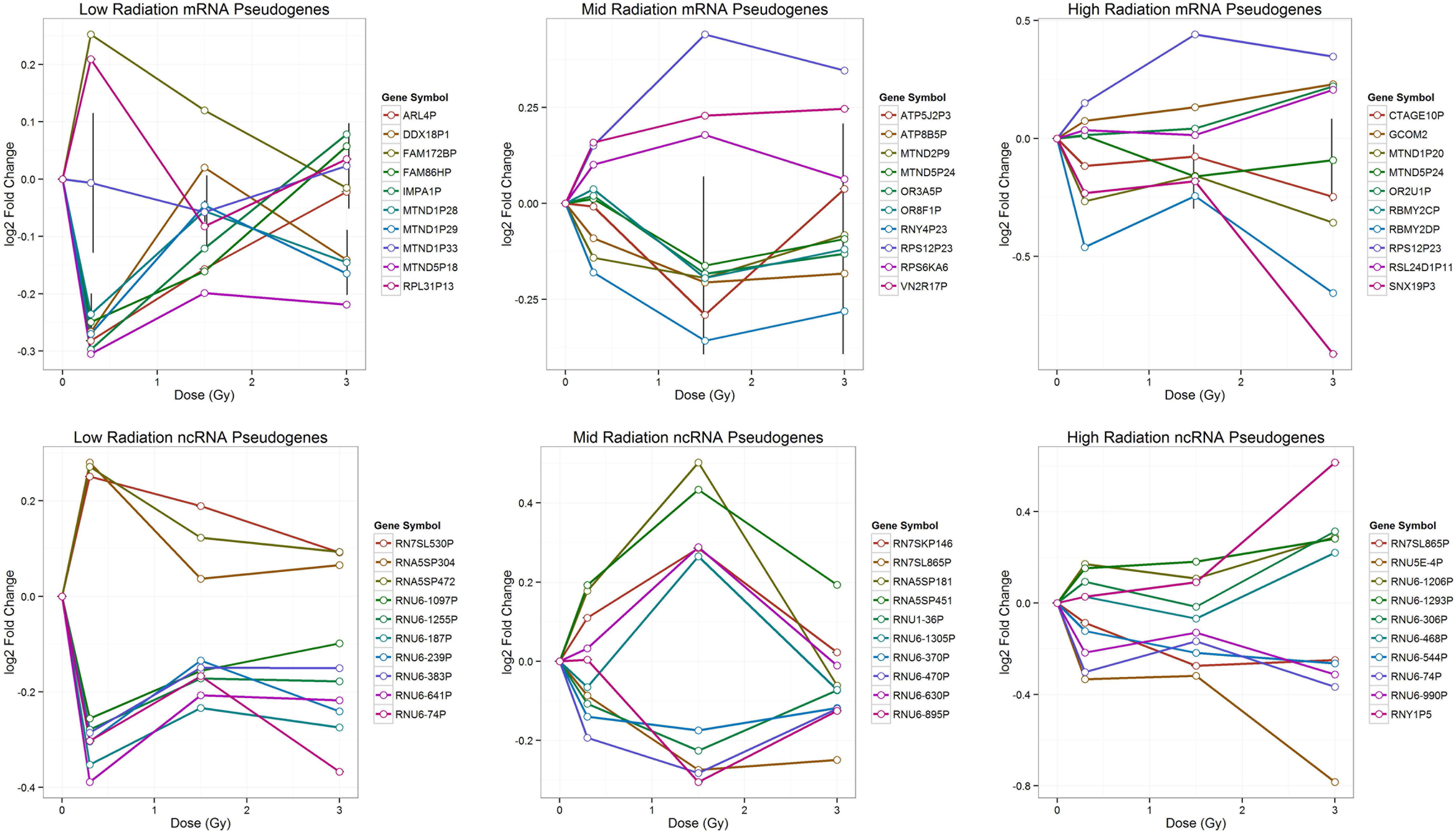
mRNA and ncRNA pseudogene expression patterns for differentially expressed mRNA (top) and ncRNA (bottom) pseudogene transcripts at 0.3 Gy (left), 1.5 Gy (center) and 3.0 Gy (right).

The majority of the down-regulated pseudogenes were pseudogenes for the mitochondrially encoded NADH dehydrogenase 1 (*MTND1*) or were related to ATP activity, suggesting the down-regulation of mitochondrial specific pseudogenes which may act as sponges for regulatory elements for their parent genes. Other down-regulated pseudogenes were related to ribosomal processes. In fact, all of the top ncRNA pseudogenes (both up- and down-regulated) fell into this category.

### Categorical enrichments

Significantly enriched GO::BP categories with a minimum gene list of five (Tables 6-8) showed a molecular response to detection of chemical stimulus that was independent of the dosage. This agreed with prior studies that showed radiation exposure induced olfactory sensations and reduced smell acuity (76–79). At 1.5 Gy, metabolic processes were enriched (Table 7). Previous work showed that global changes in the metabolome occurred with radiation exposure (100), specifically with DNA and RNA metabolism (101–105). The highest dosage, 3.0 Gy, was enriched for a number of cell cycle categories (Table 8), indicating the cell was likely trying to compensate for a large number of DNA damage events and shut down the cellular mechanisms for mitotic reproduction.

**Table 6.**
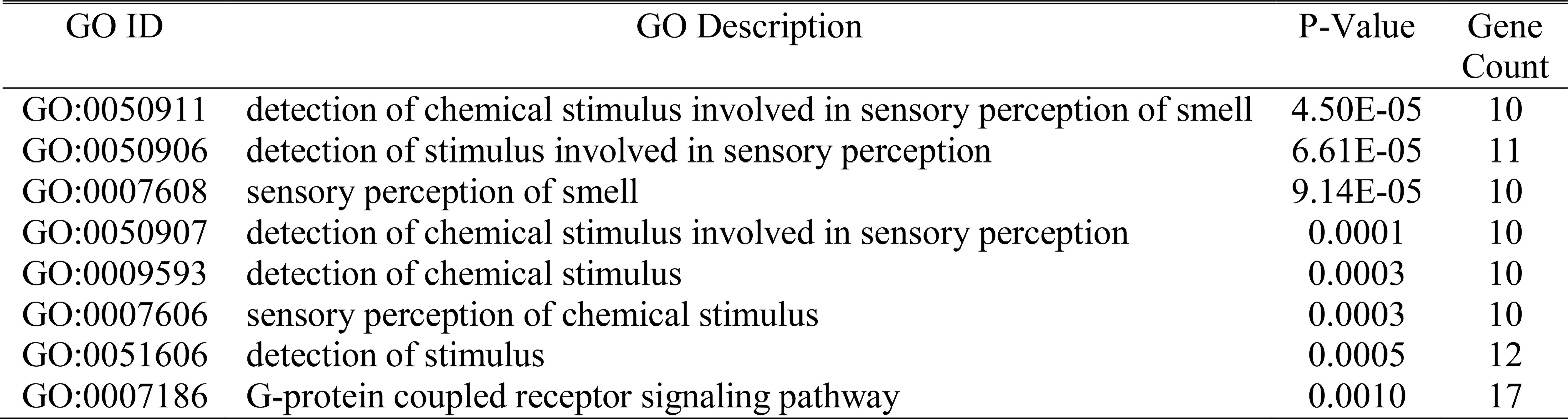
Enriched GO::BP with at least 5 genes at 0.3 Gy radiation.

**Table 7.**
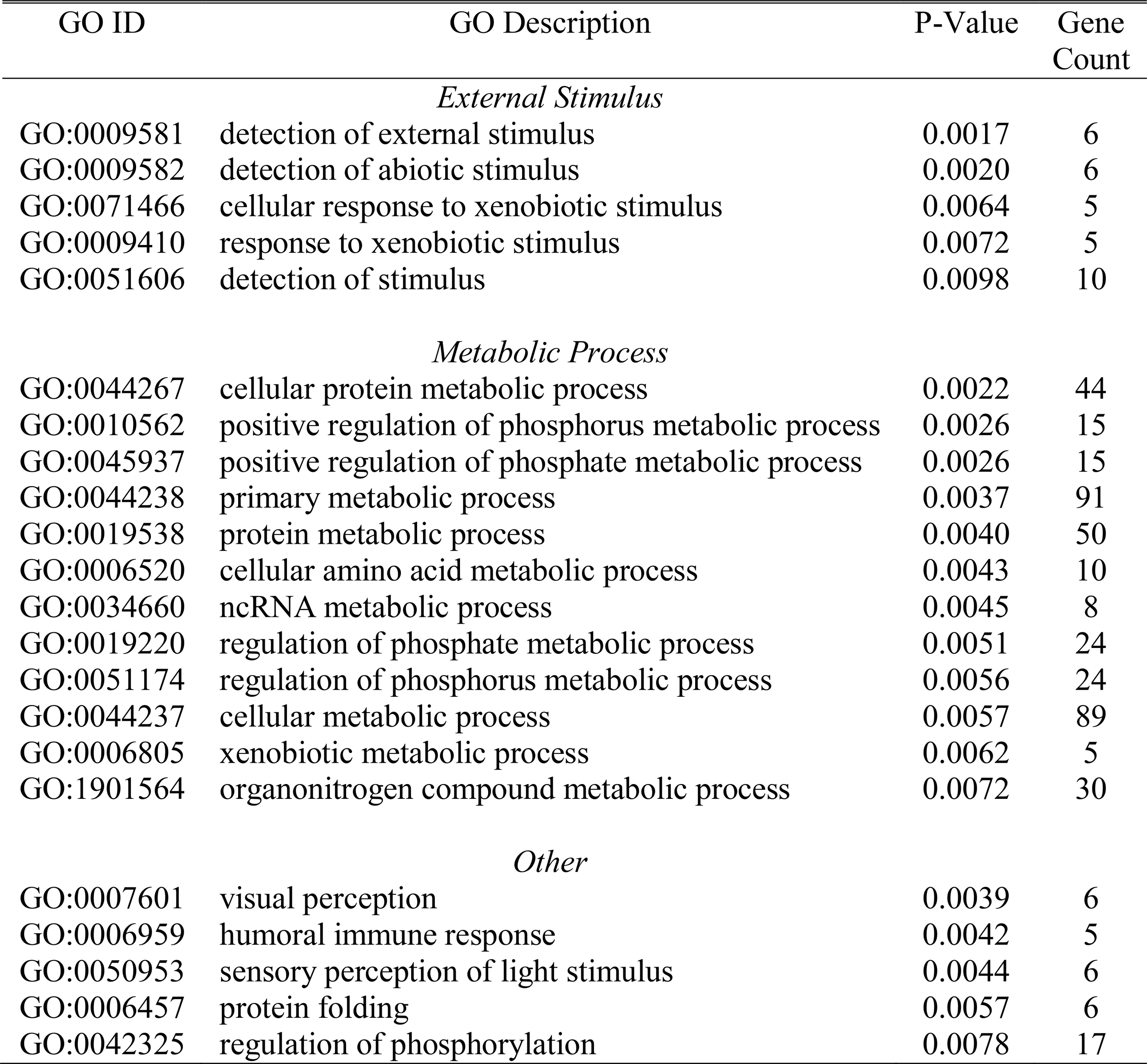
Enriched GO::BP with at least 5 genes at 1.5 Gy radiation.

**Table 8.**
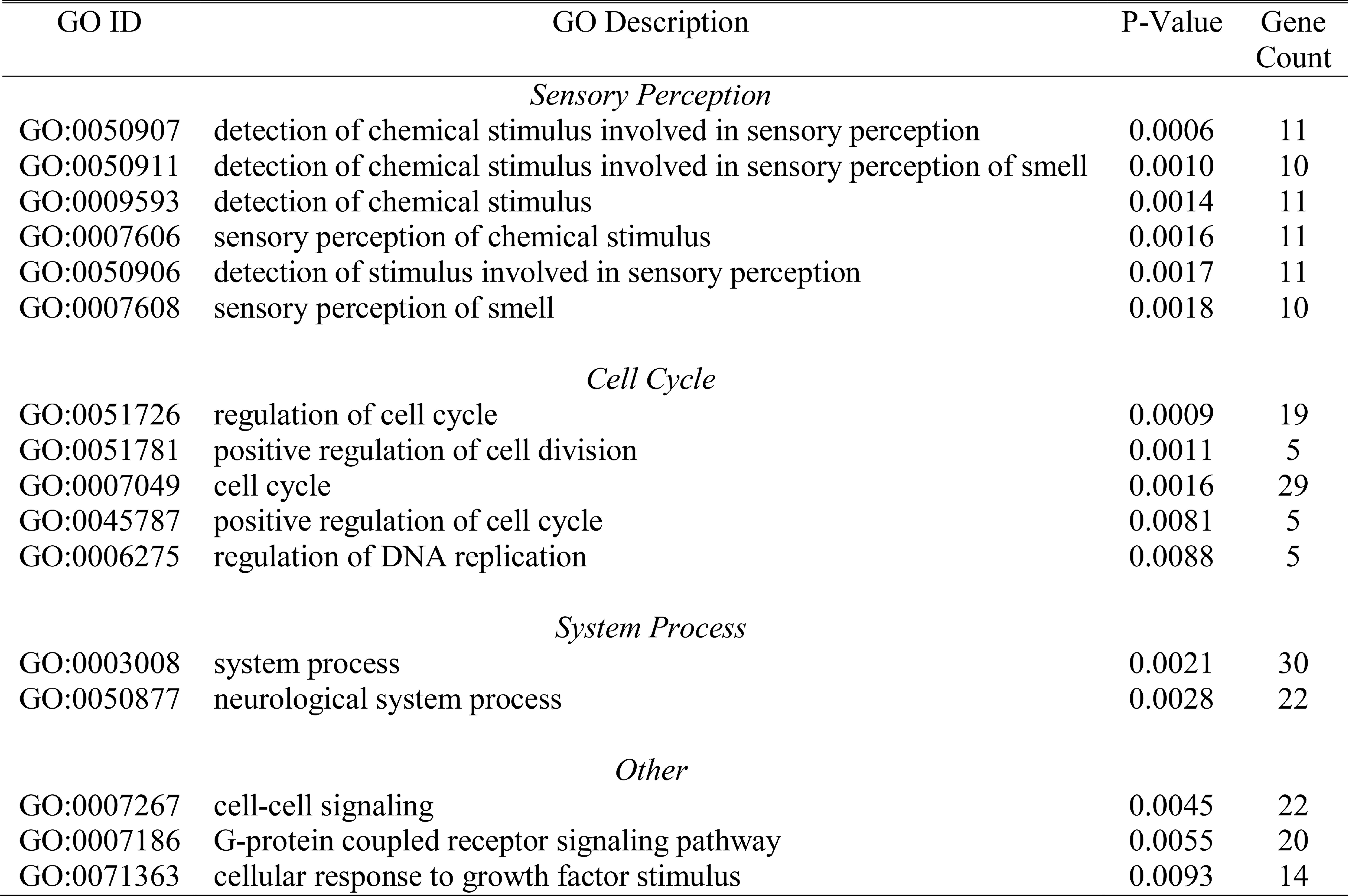
Enriched GO::BP with at least 5 genes at 3.0 Gy radiation.

Fig 7-9 highlight the significantly enriched GO::BP categories detected regardless of the number of genes in each list. Additional categories showed an up-regulation of protein transport and secretion at 0.3 Gy; increased immune response at 3.0 Gy; decreased cAMP biosynthesis and metabolism at 1.5 Gy; and decreased T-cell migration at 1.5 Gy.

**Fig 7.**
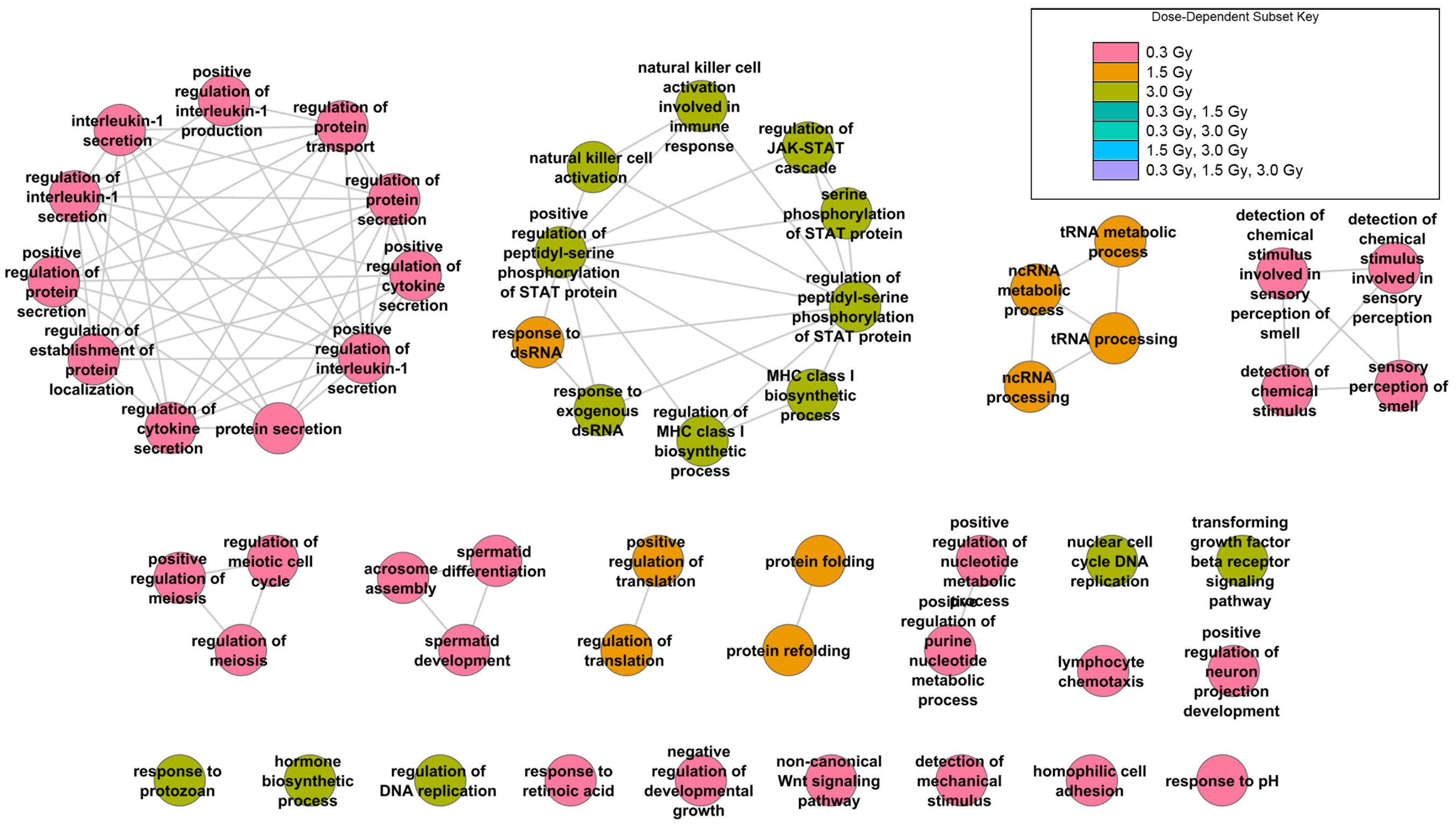
Enriched GO:BP results from categoryCompare for up-regulated DEGs at 1 hr post-exposure.

**Fig 8.**
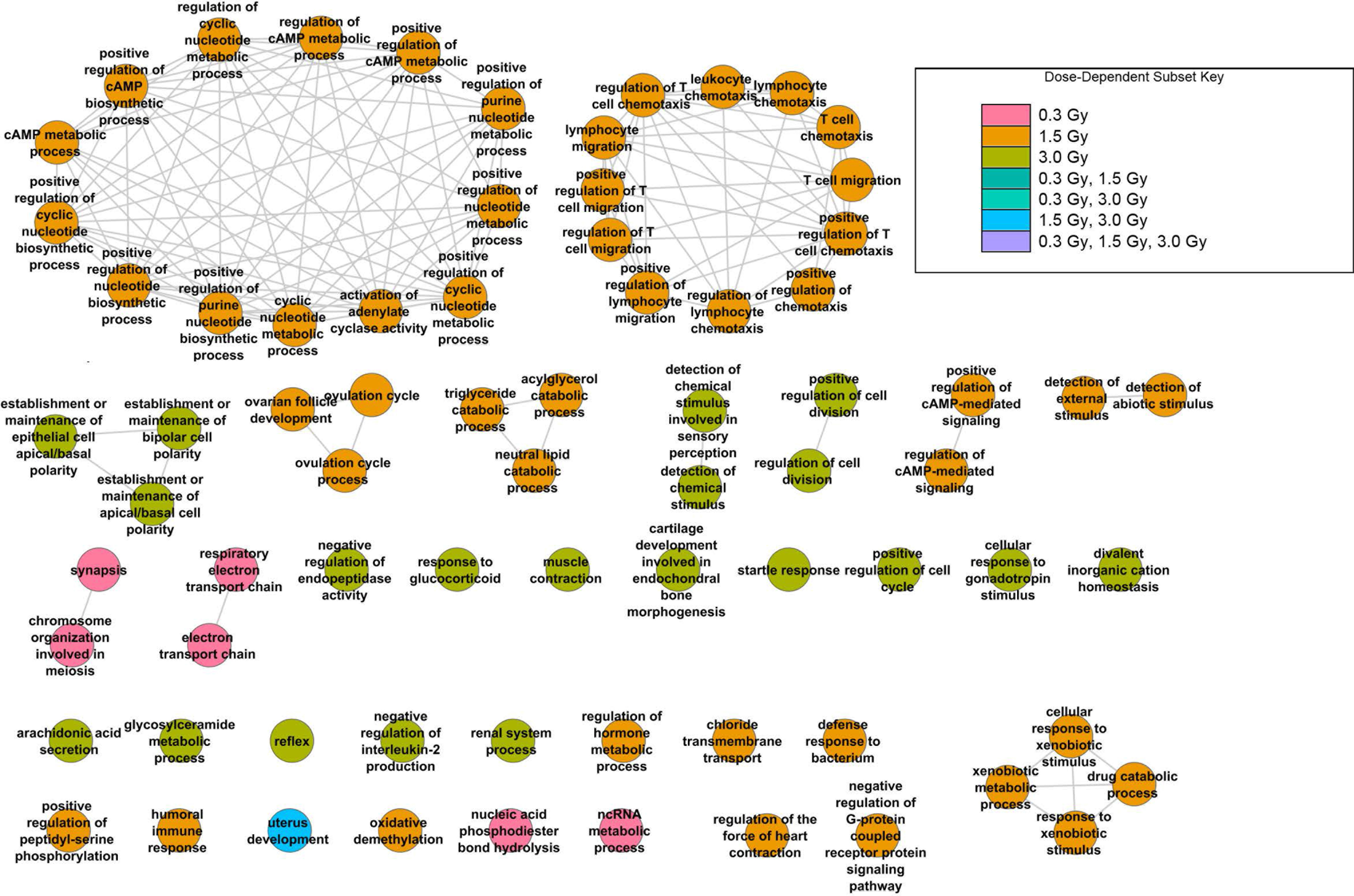
Enriched GO:BP results from categoryCompare for down-regulated DEGs at 1 hr post-exposure.

**Fig 9.**
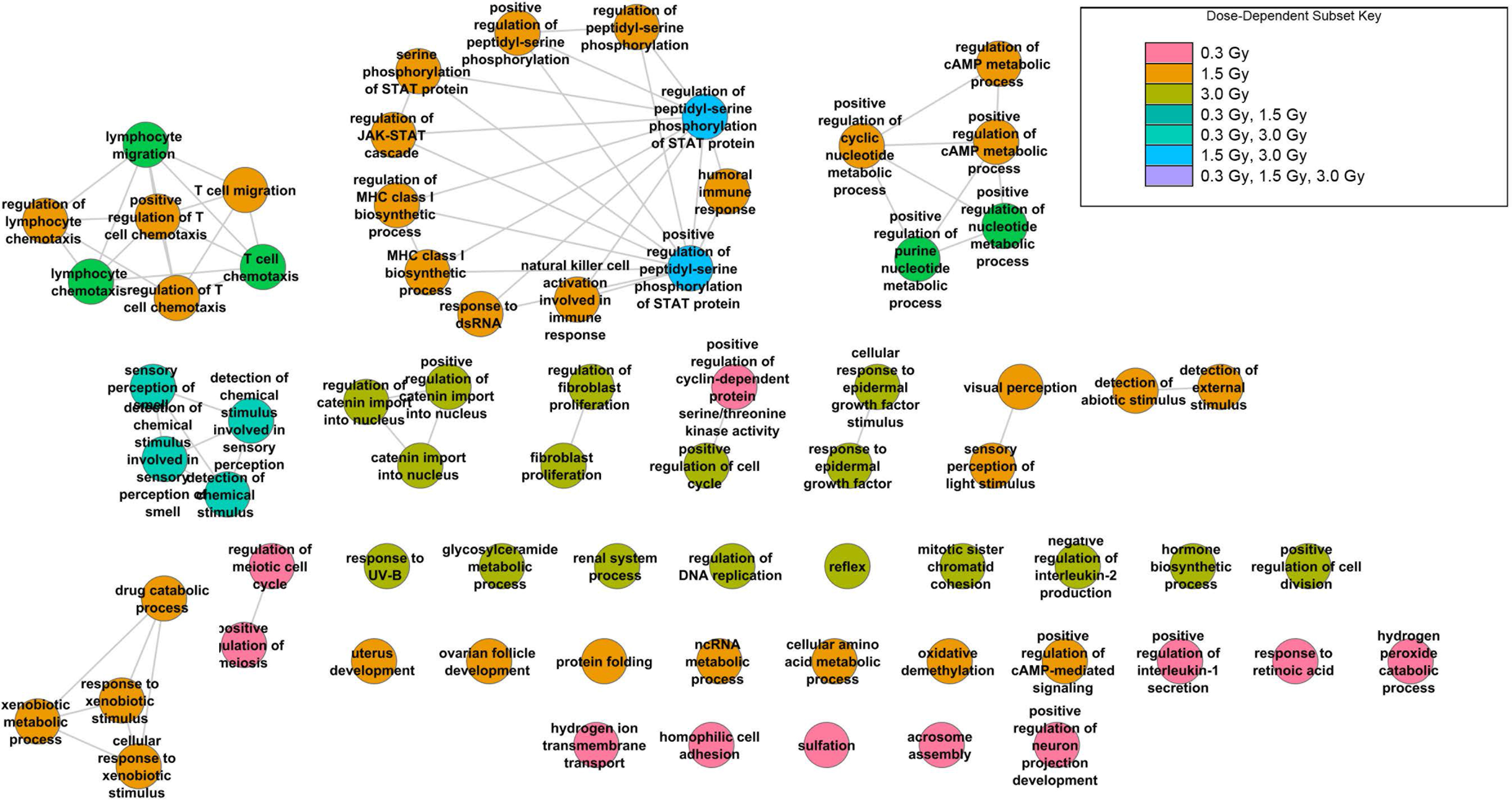
Enriched GO:BP results from categoryCompare for up-and/or down-regulated DEGs at 1 hr post-exposure.

Enriched KEGG metabolic pathways are shown in Fig 10. Among the enriched pathways were oxidative phosphorylation which, along with mitochondrial electron transport, is increased with ionizing radiation exposure (106–109). In addition, the ribosome pathway was enriched at 3.0 Gy. This was consistent with an increase in small RNA molecules responsible for rRNA maturation (Table 5) and previous results (110, 111).

**Fig 10.**
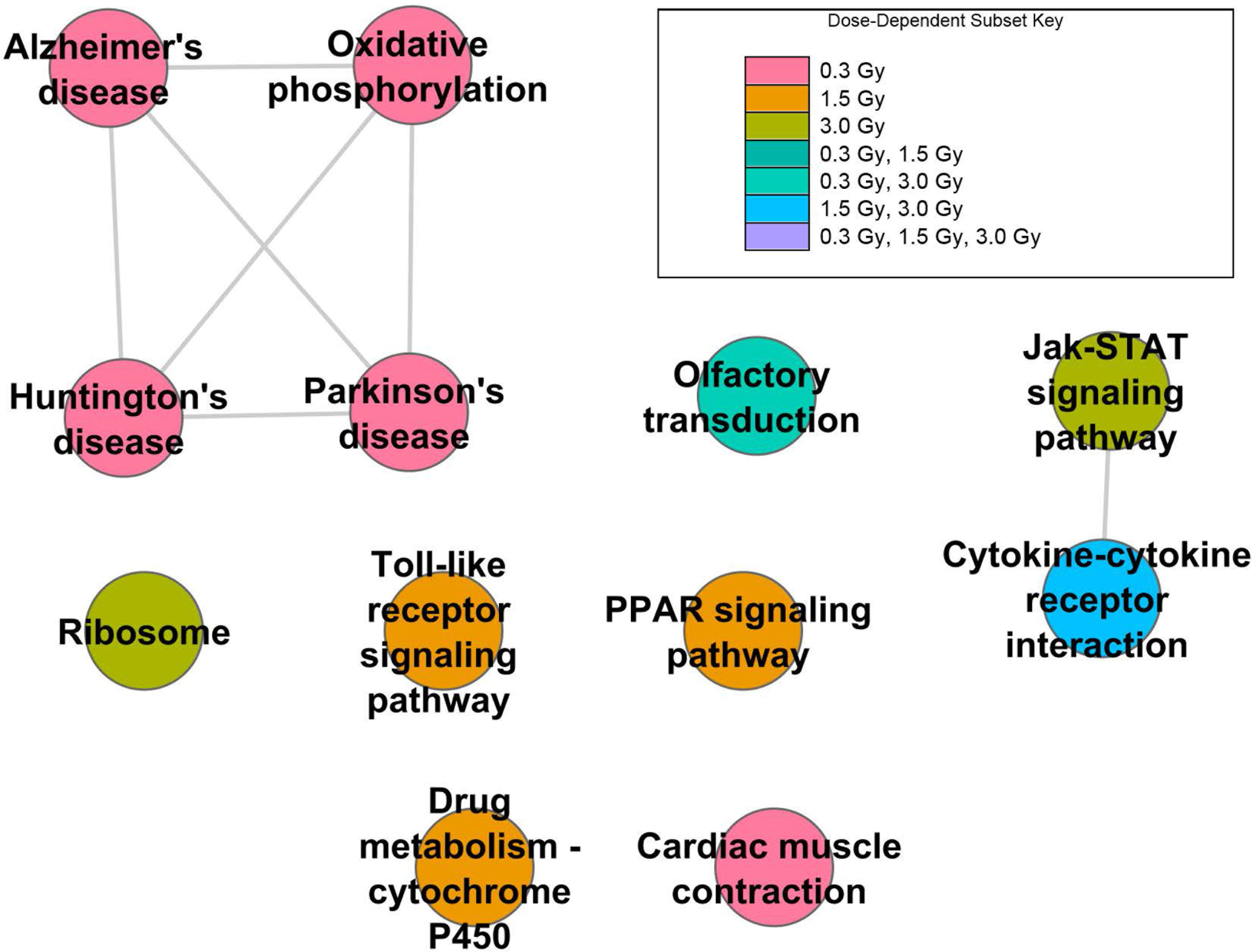
Enriched KEGG results from categoryCompare for up-and/or down-regulated DEGs at 1 hr post-exposure.

### Dose-dependent responses

Examination of the Entrez gene identifiers mapped by the differentially expressed probesets yielded 182 transcripts uniquely differentially expressed at 0.3 Gy, 161 uniquely differentially expressed at 1.5 Gy, and 223 uniquely differentially expressed at 3.0 Gy, indicating the majority of differentially expressed transcripts were dose-dependent (Fig 2). The top 10 significant dose-dependent, differentially expressed transcripts are shown in Table 9 for each of the three radiation doses.

**Table 9.**
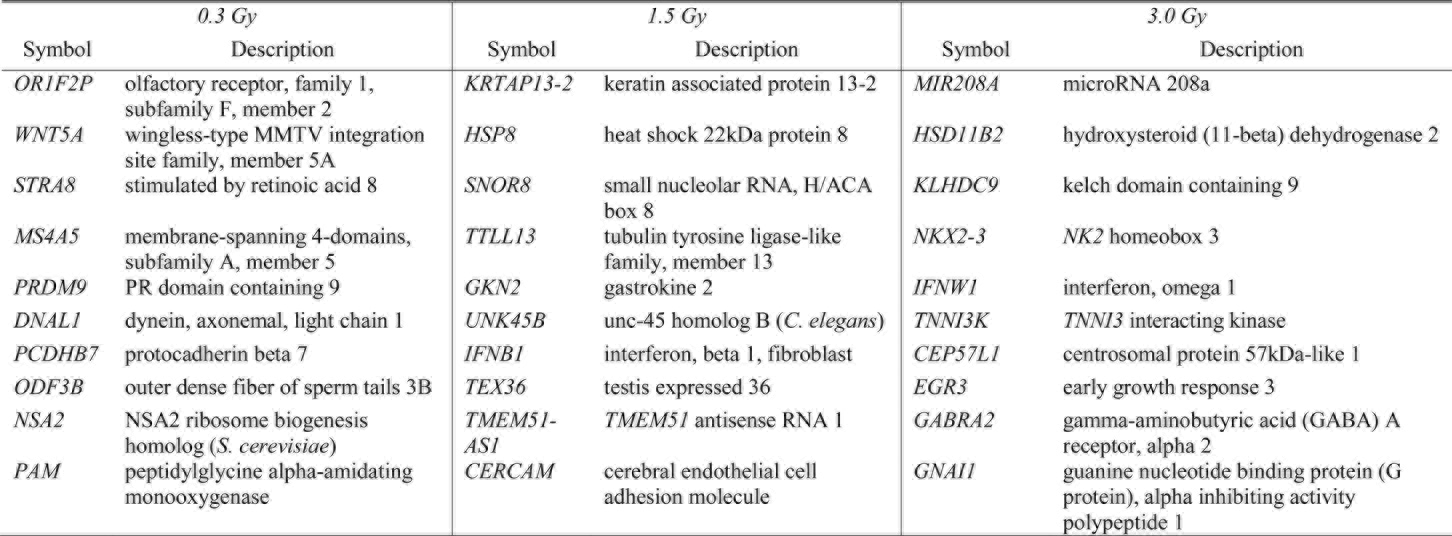
Top 10 differentially expressed dose-specific transcripts.

DAVID (112) GO::BP enrichment analysis of genes differentially expressed at a specific radiation level was performed with a minimum gene count of 6 and a significance cutoff of p ≥ 0.05 (Table 10). Since this was a reduced gene list, the number of enriched categories was smaller than that reported for all differentially expressed transcripts at each dosage. The results suggest that cell adhesion and signaling were affected at low-dose radiation (0.3 Gy), consistent with prior studies (113, 114). At the same time, an intermediate dose of radiation (1.5 Gy) elicited a response from transcripts involved in sensory perception. A higher level of radiation (3.0 Gy) more directly affected transcription of genes involved in cell cycle regulation and cellular homeostasis, which are more traditionally observed responses in radiation exposure events.

**Table 10.**
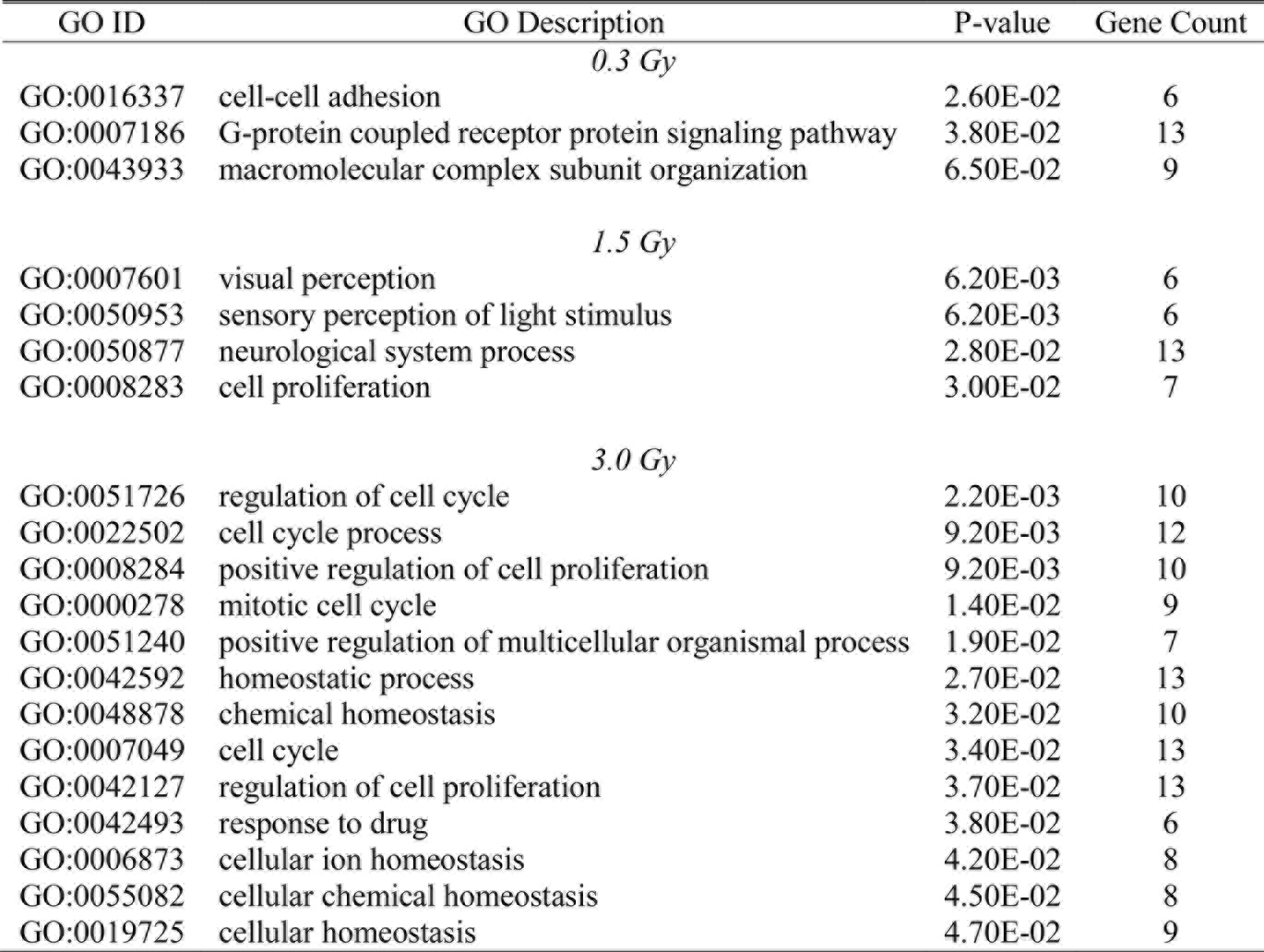
Enriched GO::BP for dose-dependent transcriptional responses.

### Sensitive and specific transcriptional biomarkers

In general, differentially expressed transcripts detected within microarray experiments yield biomarkers sensitive to a specific condition when all other factors are reasonably controlled. However, in order for a biomarker to be useful in terms of a diagnostic test, it must also be specific to the condition being tested. For example, within our dataset, *PCNA* (proliferating cell nuclear antigen) was highly differentially expressed at both 1.5 Gy and 3.0 Gy, but has been determined to be differentially regulated under a large number of conditions.

Using the methodology for determining sensitive and specific transcriptional biomarkers outlined in Materials and methods, a total of 23 genes at 0.3 Gy were considered for possible up-regulated biomarkers (Table 11), while 20 genes at 1.5 Gy (Table 12) and 23 genes at 3.0 Gy were considered (Table 13). Only one of these genes, *KDELC1* (KDEL (Lys-Asp-Glu-Leu) containing 1) was found to be differentially expressed at all three radiation levels, while *MRPS30* (mitochondrial ribosomal protein S30), *RARS* (arginyl-tRNA synthetase), and *HEXIM1* (hexamethylene bis-acetamide inducible 1) were found to be differentially expressed at both 1.5 Gy and 3.0 Gy. For each of the datasets of possible transcriptional biomarkers, we then looked at their pairwise occurrence within GEO DataSets. From this analysis, we focused on the three potential transcriptional biomarkers with the fewest pairwise occurrences within the list (Tables 14-16). For 0.3 Gy, this yielded *PRDM9* (PR domain containing 9), *CHST4* (carbohydrate (N-acetylglucosamine 6-O) sulfotransferase 4), and *SLC26A10* (solute carrier family 26, member 10). The reduced transcriptional biomarker set at 1.5 Gy included *RPH* (retinal pigment epithelium-derived rhodopsin homolog), *CCDC96* (coiled-coil domain containing 96), *WDYHV1* (WDYHV motif containing 1), and *IFNA16* (interferon, alpha 16) while those detected at 3.0 Gy included *CWC15* (*CWC15* spliceosome-associated protein), *CHCHD7* (coiled-coil-helix-coiled-helix domain containing 7), and *DNAAF2* (dynein, axonemal, assembly factor 2).

**Table 11.**
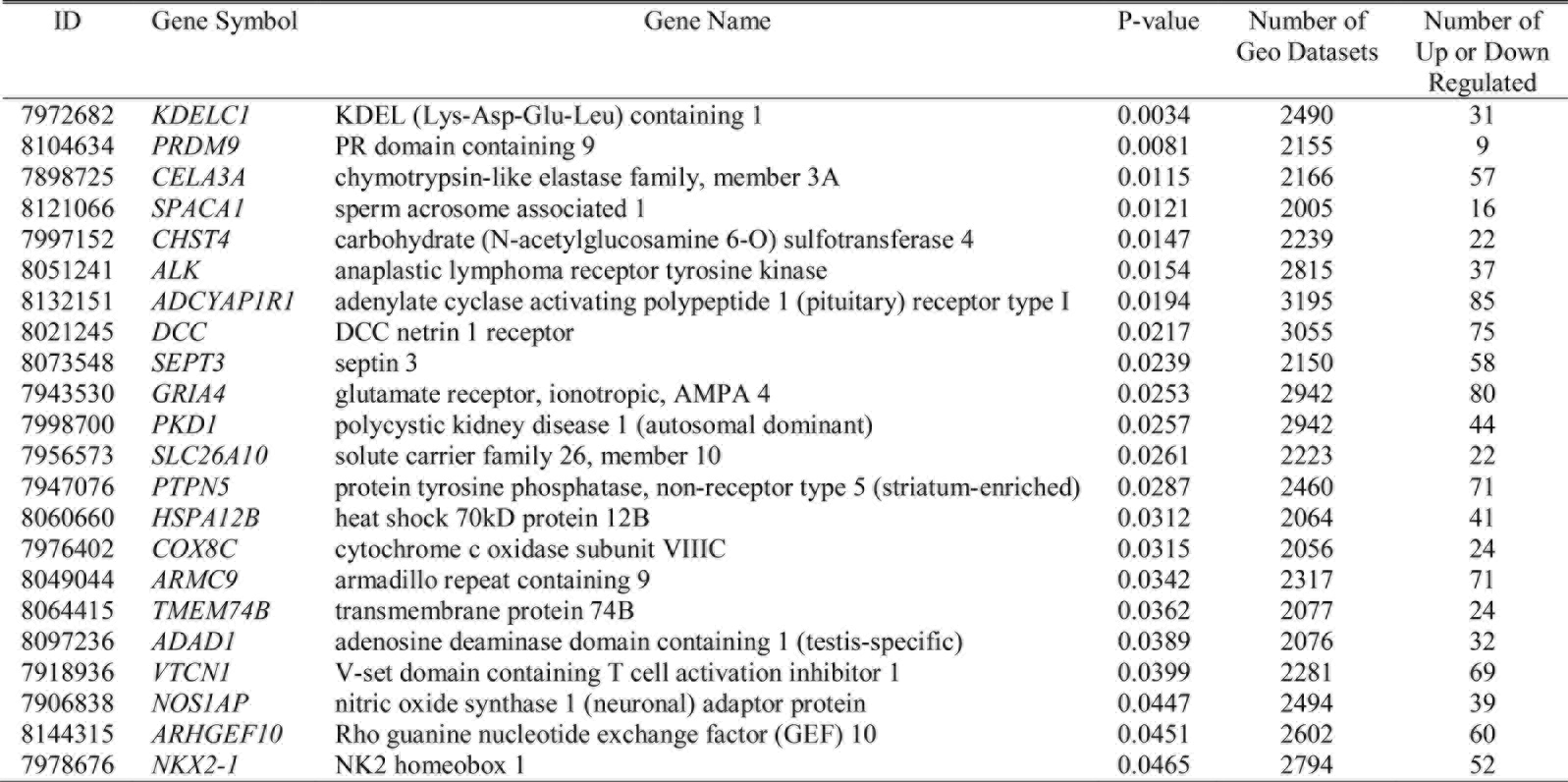
Filtered up-regulated biomarkers at 0.3 Gy radiation.

**Table 12.**
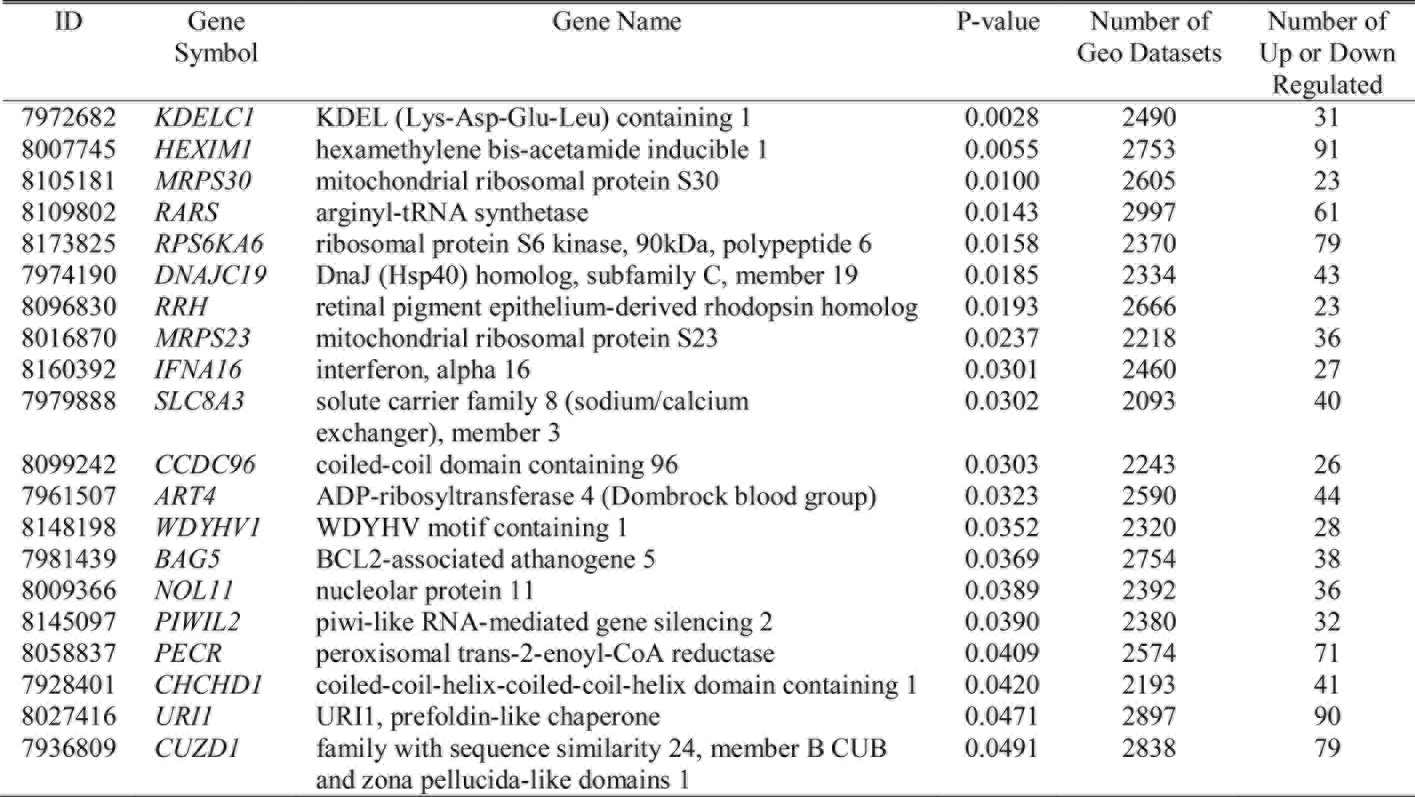
Filtered up-regulated biomarkers at 1.5 Gy radiation.

**Table 13.**
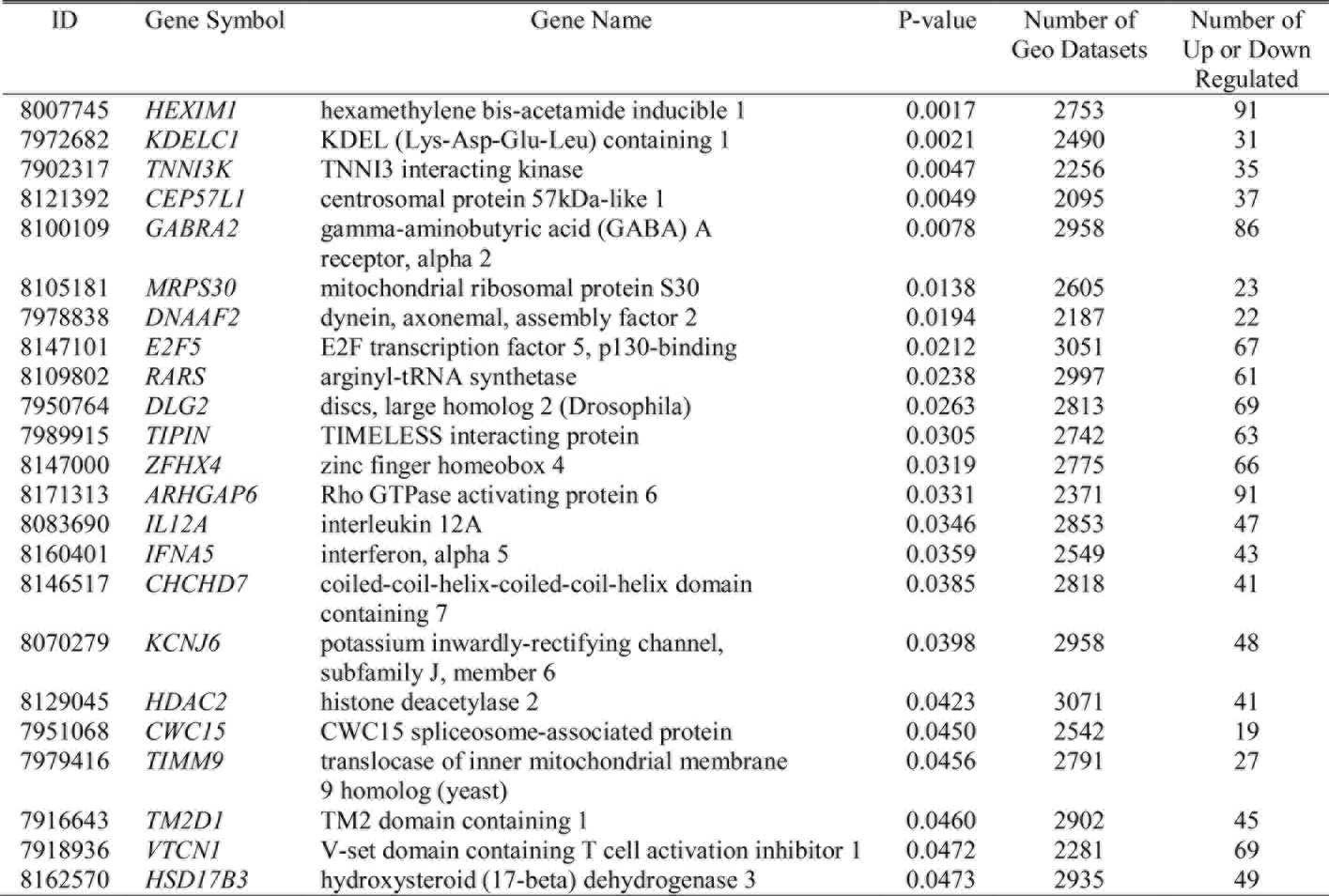
Filtered up-regulated biomarkers at 3.0 Gy radiation.

**Table 14.**
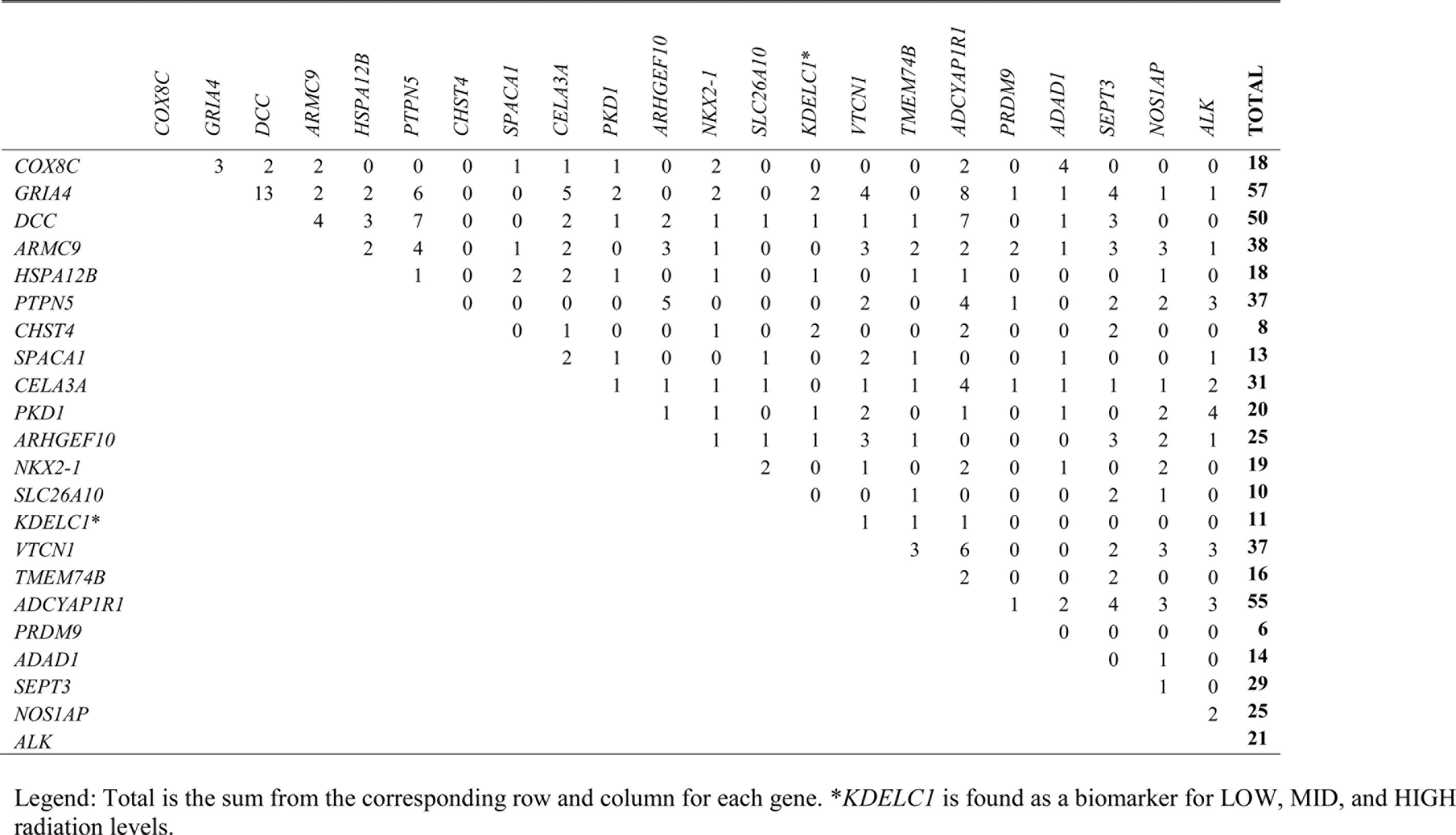
Number of shared GEO DataSets where both genes are differentially expressed for LOW radiation biomarkers.

**Table 15.**
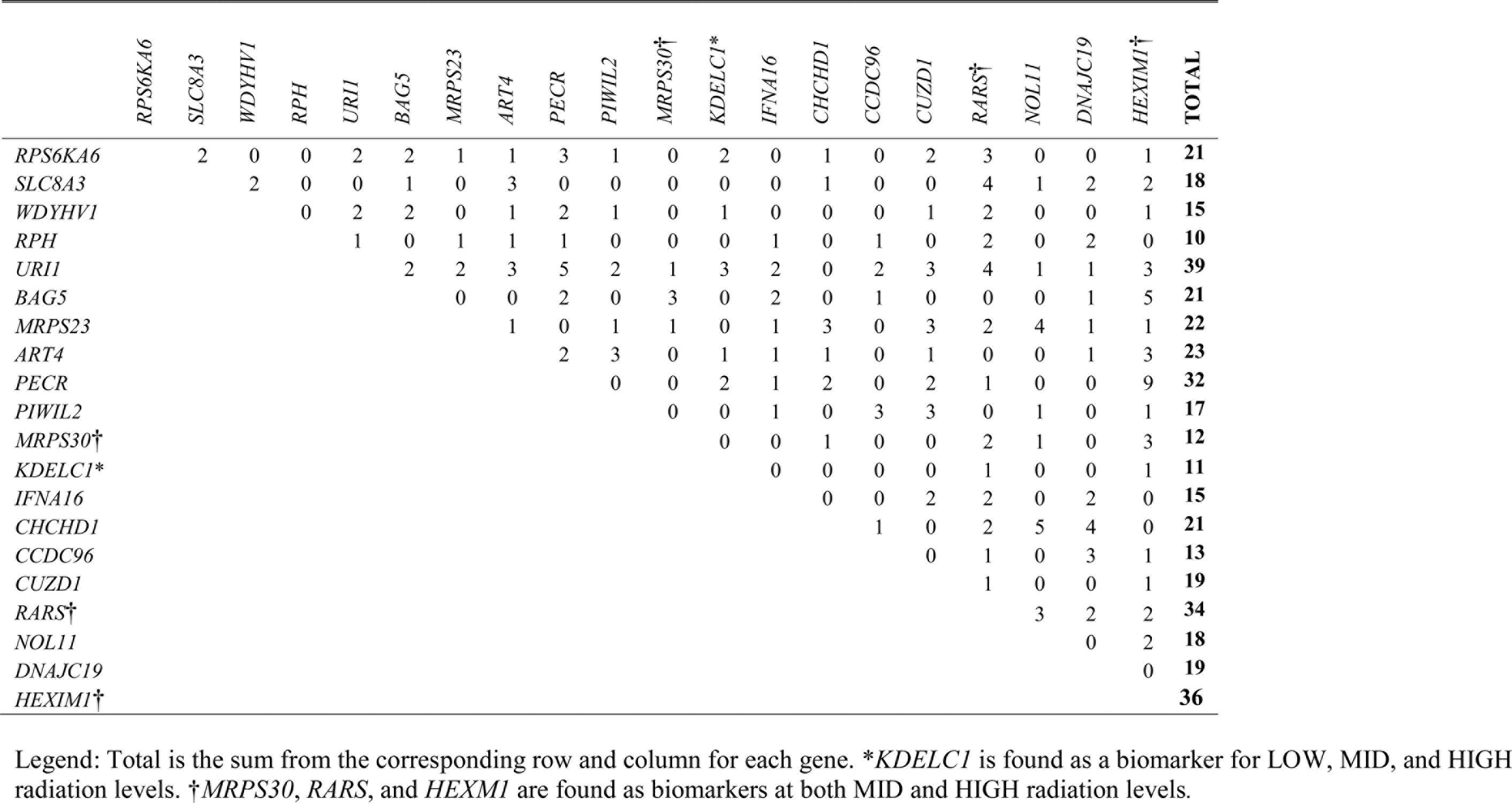
Number of shared GEO DataSets where both genes are differentially expressed for MID radiation biomarkers.

**Table 16.**
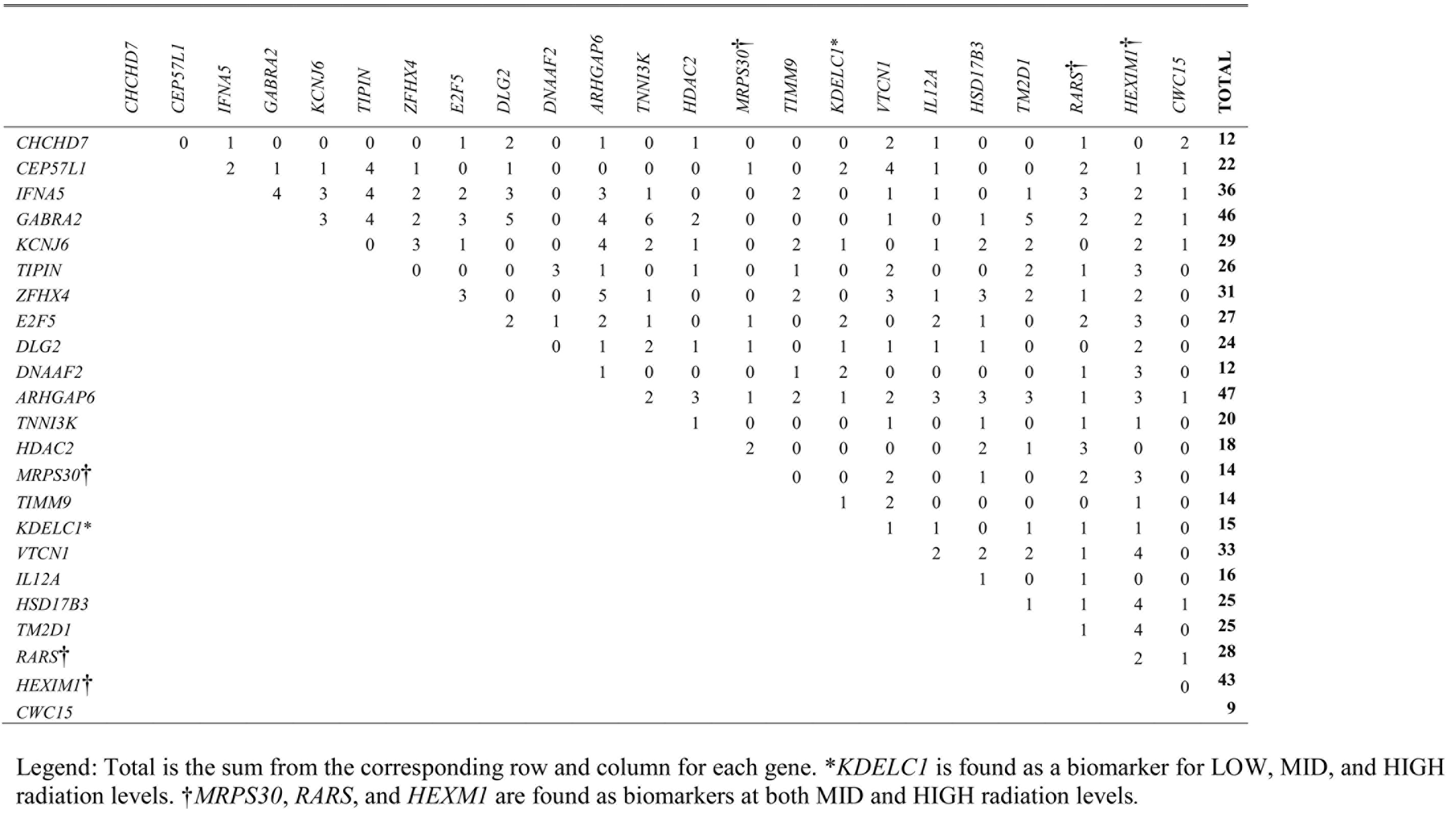
Number of shared GEO DataSets where both genes are differentially expressed for HIGH radiation biomarkers.

## Discussion

Our analysis of the transcriptional changes occurring immediately following ionizing radiation at 0.3 Gy, 1.5 Gy, and 3.0 Gy indicated a number of transcripts were significantly differentially expressed at the ncRNA, pseudogene, miRNA, and mRNA levels. Nearly all of the ncRNAs and pseudogenes with a known function or homolog that were regulated were involved in rRNA maturation or mitochondrial specific genes, indicating that radiation exposure affected the overall translation process as well as production of cellular energy. Not surprisingly, at the miRNA level, most of the regulated miRNAs were involved in regulating genes associated with cell cycle or tumor processes. The resulting responses at the mRNA level indicated both a general radiation response that was highly enriched for genes involved in sensory perception as well as dose-dependent responses where 1.5 Gy induced responses to genes involved in metabolic processes and 3.0 Gy was enriched for genes involved in the cell cycle and cellular signaling.

As a result of this study, we were also able to elucidate additional transcriptional biomarkers for general radiation exposure, including *KDELC1*, *MRPS30*, *RARS*, and *HEXIM1*, as well as dose-specific markers including *PRDM9*, *CHST4*, and *SLC26A10* (0.3 Gy); *RPH*, *CCDC96*, *WDYHV1*, and *IFNA16* (1.5 Gy): and *CWC15*, *CHCHD7*, and *DNAAF2* (3.0 Gy) which were both sensitive and specific to their corresponding radiation exposure immediately (1 hr) following exposure. Many of the differentially expressed ncRNAs and genes (including the elucidated biomarkers *MRPS30* and *CHCD7*) were either localized within the mitochondria or have mitochondrial functions, consistent with previous findings that ionizing radiation increased mitochondrial oxidative stress (106, 107, 115, 116) and altered mitochondrial function (107, 117–120). Many others, including the transcriptional biomarkers *KDELC1*, *RARS*, *HEXIM1*, *PRDM9*, and *CWC15* played a role in ribosomal maturation and activity, which has been shown as key in survival after radiation damage (110, 121, 122).

Detection of globally enriched biological processes and pathways provides partial insight into how the differentially expressed transcripts function. However, in some cases, the known function of a particular gene in radiation exposure is not clearly evident. As one example, *KDELC1* is differentially up-regulated at all three doses. However, its known function as a luminal protein allows resident proteins to escape the endoplasmic reticulum, which does not yield clear insight into its function in radiation exposure. In this particular case, *KDELC1* has been shown to form a sense-antisense gene pair (SAGP) on chromosome 13 with basic immunoglobulin-like variable motif (BIVM). SAGPs are transcribed together in pairs in opposite direction on complementary DNA strands. This particular SAGP is part of a group of 12 whose differential expression has been determined to be highly predictive of breast cancer survivability (123). Further analysis of BIVM shows that it often forms a read-through transcript with excision repair cross-complementary rodent repair deficiency, complementation group 5 (*ERCC5*) (124). The BIVM-ERCC5 functions in single-stranded DNA binding, nucleotide excision repair, nucleic acid phosphodiester bond hydrolysis, and endonuclease activity, all of which may be in direct response to radiation exposure. Therefore, *KDELC1* differential expression could directly signal a high expression of BIVM-ERCC5.

Our transcriptional profile has led to a number of elucidated transcriptional biomarkers that have not been previously implicated in radiation exposure. Many of these have roles in related processes and show a specificity and sensitivity that make them prime candidates for inclusion on diagnostic platforms. For others, functional annotation may not be complete, making their use as transcriptional biomarkers subject to additional scrutiny. As the *KDELC1* example illustrates, this functionality in radiation exposure may be an indirect association, allowing for a novel transcriptional biomarker resulting from the complex interaction among multiple transcripts. The results of our study using microarray technology have yielded a number of transcriptional biomarkers that can be flagged for further transcriptomic analysis of expression, including alternative 5’ and 3’ UTR usage and alternative splicing using current RNA-Seq technologies.

## Acknowledgements

The authors wish to thank members of the University of Louisville Research Group for Diagnosing and Mitigating Human Exposure to Radiation Using Micro-Nanotechnology and members of the Kentucky Biomedical Research Infrastructure Network Bioinformatics Core for critical feedback. We wish to thank the anonymous reviewers for their time and feedback as well.

